# Effective concentration of marine nanoflagellates with a microfluidic device

**DOI:** 10.1101/2023.04.03.534374

**Authors:** Bryan R. Hamilton, Kristen R. Hunter-Cevera

## Abstract

Protist cells are typically manipulated through either centrifugation or membrane filtration, which can damage these fragile cell types. Use of microfluidic devices could greatly aid in the separation and concentration of protist cells with significantly less damage. Recent developments have enabled passive cell separation and consequent concentration based only on cell size. We utilize these advances to show that a passive spiral microfluidic device can effectively concentrate marine nanoflagellates within the 3-20 micron size range without harm to cells, while reducing background bacteria levels. The ability to concentrate these cell types appears only dependent on cell size, despite complicated cell surface geometries and motility. We anticipate that this approach will greatly aid researchers who require an ability to manipulate fragile cell types as well as reduce bacteria concentrations for experimental setups and cell isolation.

## Introduction

Nanoflagellates are small protist cells (< 20 microns) that play a key role in marine biogeochemical cycling and oceanic food webs (Azam et al. 1983; Li et al. 2021; Corradino and Schnetzer 2022). These cells are major consumers of marine bacteria and other members of the picoplankton, serving as an important trophic link and aiding in nutrient remineralization (Das and Pandey 2015; Johnke et al. 2017; Corradino 2020; Zhang et al. 2021). To study such cells in the laboratory requires general manipulation. However, these cells are inherently fragile due to a wide diversity of shapes, motilities, structures (including extracellular spines), and other apparatuses that either aid in feeding or defense (Arndt et al. 2000; van Tol et al. 2012). Common mechanisms to concentrate them, such as membrane filtration or centrifugation, typically result in cellular damage (Bloem et al. 1986; Andersen 2005).

The inability to safely manipulate, concentrate, and/or sort these cells has affected how they can be studied and used in laboratory experiments and downstream applications. For example, nanoflagellate ingestion rate experiments are typically constrained by the maximum density of nanoflagellate cells that can be achieved in culture with underlying background bacteria populations (Apple et al. 2001; Guillou et al. 2001; Christaki et al. 2002). Processes such as prey ingestion though show density dependence (Altermatt et al. 2015; Yan et al. 2018), such that the ability to concentrate grazers while lowering background bacteria levels would enable more precise control over experimental designs and better representation of conditions found in the marine environment.

The ability to concentrate and sort nanoflagellates would also aid in isolation attempts. Most of our knowledge pertaining to nanoflagellate genomics and their ecological role in marine environments is limited to only a handful of easily cultivable examples (Sherr et al. 2007). These organisms are often not the most abundant nor relevant members in the environment (del Campo et al. 2013). The ability to size separate nanoflagellates from other cell types in marine samples and enrichment cultures could be a valuable tool for successful isolation and long term culture attempts.

In recent years, there has been an increasing interest in the use of microfluidic devices to perform cell separation tasks (Shields IV et al. 2015, Zhou et al. 2019). Cell separation methods can be categorized as active, passive, or a combination of the two (Fan et al. 2014; Rostami et al. 2020). Active methods require external forces such as magnetic, acoustic, or electric power to separate cells in a sample. Passive methods do not rely on external forces, but instead use channel geometry and hydrodynamic forces to separate cells or other particles based on their physical properties (Russom et al. 2009; Guan et al. 2013). Many researchers are exploring passive cell separation methods due to their simple design, low cost, label-free, and straightforward operation (Zhou et al. 2019; Pensold and Zimmer-Bensch 2020).

Spiral micro-channel designs are a passive method that have been used for a wide variety of applications. Particles in a spiral micro-channel are mainly affected by two forces: inertial lift force and Dean drag force (Nivedita et al. 2017; Condina et al. 2019). Depending on the flow rate and particle size, these forces act to position particles at an equilibrium position along channel height and width. Particles are sorted or concentrated by designing different outlets specific to that position. For more information, we refer the reader to Lee and Yao (2018) for a detailed explanation of the forces that act on particles within a spiral channel.

Successful sorting of different sized particles in the range of 5-30 microns has been shown for uniform particles (spherical beads) and for irregular shaped particles (Guan et al. 2013; Roth et al. 2018). Studies have also demonstrated the usefulness of spiral microfluidic devices for the concentration and separation of different cell types and for separation of cells from biological fluids (Bankó et al. 2019; Liu et al 2021). For example, with a spiral microfluidic device, Xiang et al. (2019) separated rare tumor cells from red and white blood cells in whole blood samples. Separation of target cells from smaller bacteria cells (which are not typically affected by the forces within a spiral device) is also an important application. Hou et al. (2015) separated low abundance bacteria from host cells in whole blood samples to identify pathogenic bacteria, and Condina et al. (2019) created a device to separate beer spoilage bacteria from yeast cells in beer samples, which could then be used in downstream mass spectrometry to quickly identify contaminants. Both of these examples eliminated the need for a time consuming culturing step to determine if bacteria were present in samples.

While such microfluidic spiral devices have been able to size select mammalian and other eukaryotic cells, to our knowledge, they have not yet been utilized for non-destructive sorting and concentration of aquatic protists cells. In this study, we explore the use of a microfluidic spiral device to sort and concentrate four different nanoflagellate species, while reducing the concentration of background bacteria. We find that a spiral device can successfully and non-destructively concentrate these cells down to a size of 3 microns, enabling them to be used in downstream experiments or isolations. Furthermore, each nanoflagellate species used in these experiments has a different average cell size and morphology and is motile. These organisms provide interesting test cases for the evaluation of these devices to concentrate particles with complicated cell surface geometries, flagella extensions, and motility.

## Materials and Procedures

### Device fabrication

Our microfluidic device is based on the rectangular spiral device of Guan et al (2013). The spiral consists of eight loops with a single inlet and four outlet channels. The radius increases from 8 mm at the inner loop to 24 mm at the outer loop of the spiral. The main channel is 600 μm wide and 80 μm deep, with four outlets splitting off from the main channel at the end of the outermost loop (Fig. 1). Each outlet channel has a width of 150 μm. The mold for this pattern was drawn with Blender 3D creation software (v 2.93.6), and was 3D printed with a Form3B SLA printer (Formlabs), with clear resin. Following the print, the mold was washed in 100% isopropanol for 10 minutes and post cured for 30 minutes with purple-blue light (wv = 405 nm) in a Form Cure machine (Formlabs). The mold was then heated at 65 °C for 4-5 hours in a convection oven (Toshiba).

**Figure 1.**
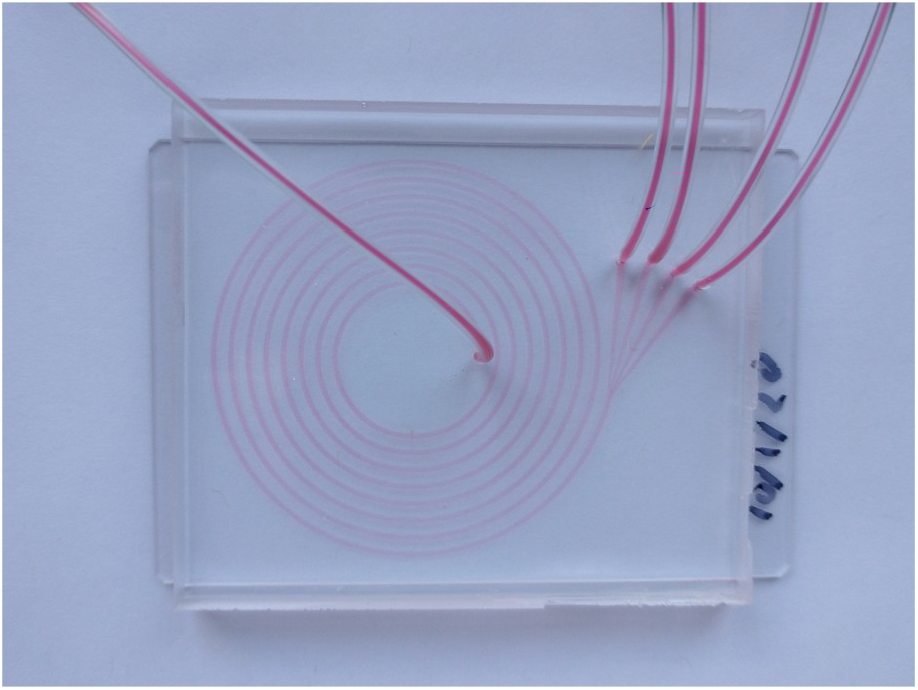
Image of spiral microfluidic device used in this study. Red food coloring highlights channel geometry and outlets leading to microcentrifuge collection tubes.

Polydimethylsiloxane (PDMS, Sylgard 184® Dow Chemicals) was prepared as a 10:1 base elastomer to curing agent mixing ratio and poured onto the cured 3D printed mold. PDMS was allowed to degas under vacuum for 30 minutes. PDMS was cured at 65 °C on a hotplate for 1-2 hours. Cured PDMS was carefully peeled off the mold and inlet and outlet holes were punched with a 1.5 mm diameter tissue punch. The PDMS and a glass slide, previously cleaned with First Contact™ Polymer (Photonic Cleaning Technologies), were plasma treated with a Plasma-Prep machine (SPI) for 20 seconds at ~50 W. The PDMS surface was immediately pressed to the glass slide for 1 minute. The device was then placed on an 80 °C hot plate for 15-30 minutes to strengthen the bond. Flexible tubing (gauge 23) was connected to each inlet and outlet hole and held in place with surface tension.

### Nanoflagellate isolation and culturing

Protist nanoflagellates were isolated from seawater collected at the Martha’s Vineyard Coastal Observatory (MVCO), located a mile off the south shore of Martha’s Vineyard Island, Massachusetts. Water was pre-filtered with a 20 μm Nytex mesh, and 5-10 mL of filtrate was dispensed into autoclaved glass test tubes. Autoclaved rice grains, as well as cultured strains of marine *Synechococcus*, were added to these tubes to encourage growth of protists and to screen for nanoflagellates that ingest *Synechococcus*. Rice grains serve as an energy source for background bacteria, which in turn provide food for the nanoflagellates. Tubes were checked for activity every few weeks by visualizing samples with a microscope. Organisms of interest were isolated using a combination of dilution series as well as manually picking individual cells with a 10 μL pipette. Isolated cultures were maintained with rice grains in filtered seawater.

Nanoflagellates were identified based on 18S rRNA sequence and existing morphology. Dense culture of isolates (~50,000 cells mL^-1^) believed to be clonal were gently spun down using a micro centrifuge and DNA was extracted using a DNeasy Microbial kit (Qiagen) following manufacturer’s instructions. The 18S rRNA gene was amplified via PCR with general 18S primers, F-566 (5’-CAGCAGCCGCGGTAATTCC-3’) and R-1200 (5’-CCCGTGTTGAGTCAAATTAAGC-3’), using an AmpliTaq Gold® DNA Polymerase kit (Applied Biosystems). Each 50 μL reaction contained 1X Buffer II, 0.2 mM DNTP, 1.5 mM MgCl_2_, 1.25 U DNA Polymerase, 0.2 μM forward and reverse primers, 37 μL of nuclease free water (Ambion), and ~15 ng of DNA template. Cycling conditions were 95 °C for 10 minutes; followed by 35 cycles of 45 seconds at 95 °C, 45 seconds at 60 °C, and 60 seconds at 72 °C; with a final extension step of 10 minutes at 72 °C.

PCR product was confirmed using gel electrophoresis, and product was cleaned following standard isopropanol wash and ethanol precipitation. Direct PCR products were sequenced at the University of Chicago Comprehensive Cancer DNA Sequencing Facility. Chromatograms were visually inspected for quality; absence of double peaks or noisy chromatograms likely indicated that cultures were clonal. Sequences were checked against NCBI Genbank database to infer most likely genus and/or species.

We utilized four resulting protist cultures: *Paraphysomonas bandaiensis* (99% identity match), *Paraphysomonas butcheri* (99% identity match), *Pseudobodo tremulans* (100% identity match), and a *Goniomonas* spp. (97% identity match) in below experiments. A bacteria control, belonging to the *Shimia* genus within the *Rhodobacteraceae* (strain 13.20C-C) was also investigated (Hamilton &. Hunter-Cevera unpubl.). This strain was grown and maintained in yeast extract-peptone-glucose media. Approximate size range for each organism was determined by light microscopy with an Axioskop 2 (Zeiss).

### Scanning electron microscopy

Nanoflagellates can vary in size and morphology, some of which cannot be detected with a light microscope. To better resolve any features, we imaged protists with scanning electron microscopy. To prepare cells for imaging, a protocol modified from Park et al. (2003) was used. Nanoflagellate cultures were fixed with 2% glutaraldehyde for 30 minutes and cells were filtered onto a 13 mm polycarbonate membrane filter with 1 μm pores (GVS). The filters were washed 3 times for 5 minutes each using 0.1 M sodium cacodylate buffer (pH 7.4) followed by a dehydration step overnight in 75% ethanol. Filters were then submerged in 100% ethanol three times for 10 minutes each before being placed in an EM CPD300 critical point dryer (Leica). Exchange cycles was set to 12. Speed settings for “CO_2_.in” and “gas-out” were set to “slow” to retain any scales that may have been present on the exterior of the cells. Samples were sputter coated with 7 nm of platinum (Leica EM MED020 Coating System) before being viewed with a Supra 40VP scanning electron microscope (Zeiss) at 3kV from secondary electron scatter.

### Experimental setup

We investigated the ability of the spiral microfluidic device to concentrate protists into four different outlet channels. We tested four different nanoflagellates (3-20 μm) with different cell morphology. We also tested the ability of the device to concentrate a bacteria species as a control. For each experiment, we utilized 5 mL Luer-Lok™ syringes (BD) and a Harvard Apparatus syringe pump (model PHD ULTRA). The spiral device was first primed with 2 mL of 0.22 μm filtered seawater before each sample to clear the channels. Approximately 5 mL of sample in a separate syringe was loaded and connected to the device via tubing. A microfluidic bubble trap (Elveflow) was fitted between the syringe and spiral device to eliminate bubbles introduced during connections. One mL of sample was allowed to run through the device prior to output collection to ensure emptying of primer wash before sample collection. Sample was pumped through the spiral device at 0.5, 1, and 2 mL/min, with a brief pause in between each speed to refill the syringe with 5 mL of fresh sample. Output from each channel was collected into 1.5 mL micro centrifuge tubes. Each run was stopped when at least one output channel tube was nearly full.

Duplicate 250 μL sub-samples were removed from each of the four outlet collection tubes. These samples, along with samples from the bulk culture, were fixed with glutaraldehyde (0.1% final concentration) and Pluronic F-68 surfactant (0.01% final concentration) for 1-1.5 hrs before freezing in liquid nitrogen. Remaining volumes in each collection tube were measured by a pipette to determine differences in volume output between the four channels. Samples from the bulk culture flask were collected just prior to the initial spiral pass and just after the final spiral pass. Each experiment was performed twice for each nanoflagellate species on two different days to account for differences in cell concentration and culture media conditions. For *Paraphysomonas bandaiensis* and *Pseudobodo tremulans*, the second experiment was performed with a physically different spiral device to confirm reproducibility of approach. Viability of cells after transiting through the device was first checked visually by monitoring movement patterns of cells with a dissecting microscope (AmScope), and second by starting new culture flasks using transited cells and monitoring successful growth after a few days.

### Flow cytometry

Frozen samples were thawed at room temperature and then stained with SYBR green (ThermoFisher) at 1X final concentration. Triton-X was also added to samples at 0.1% final concentration to help the SYBR stain penetrate cell membranes (Cheng et al. 2019). Cells were allowed to incubate in the dark for 1 hour at 4 °C. Counts of nanoflagellates and bacteria were determined by flow cytometry with an Attune NxT flow cytometer (ThermoFisher) equipped with a blue laser (488 nm). Samples were diluted with freshly filtered (0.22 μm) seawater before analysis. Both bacteria and nanoflagellates were distinguished by green fluorescence and characteristic size based on side scatter.

## Assessment

### Nanoflagellate cell morphology

*Paraphysomonas bandaiensis* is the largest nanoflagellate used in this experiment, with a cell diameter typically in the 5-10 μm range. These spherical cells are covered in siliceous plates with a spine protruding from each plate (Fig. 2A5). Two flagella of unequal size are observed protruding from the cell. *Paraphysomonas butcheri* is spherical and smaller in diameter (3-5 μm, Fig. 2B5). These cells are also covered with siliceous plates, but lack spines that are present in *P. bandaiensis* (plate morphology is typically unique to species within this genus). Two flagella of unequal size are present. *Pseudobodo tremulans* has an irregular spherical cell shape and is of similar size to *P. butcheri* (Figure 2C5). Typical cell diameter is 3-5 microns. No scales or spines are present on the cell exterior, but cells appear to be covered in an extracellular matrix. Undetermined *Goniomonas* species exhibited a slightly elongated cell shape with a diameter of 5-7 microns (Fig. 2D5). There are no noticeable scales or spines covering the cell. Two flagella extending from the cell are shorter in length relative to the body in comparison to the two *Paraphysomonas* species used in this study. We note the discrepancies between cell size reported here and the sizes indicated on the scanning electron micrographs, which would indicate that cells are much smaller. Given the shrinkage and distortion than can occur during the intense fixation required by SEM, especially for fragile cells, we utilize cells sizes determined by light microscopy (Menden-Deuer et al. 2001). We also note that background bacteria contained within the nanoflagellate cultures were often observed to have elongated and extended shapes.

**Figure 2.**
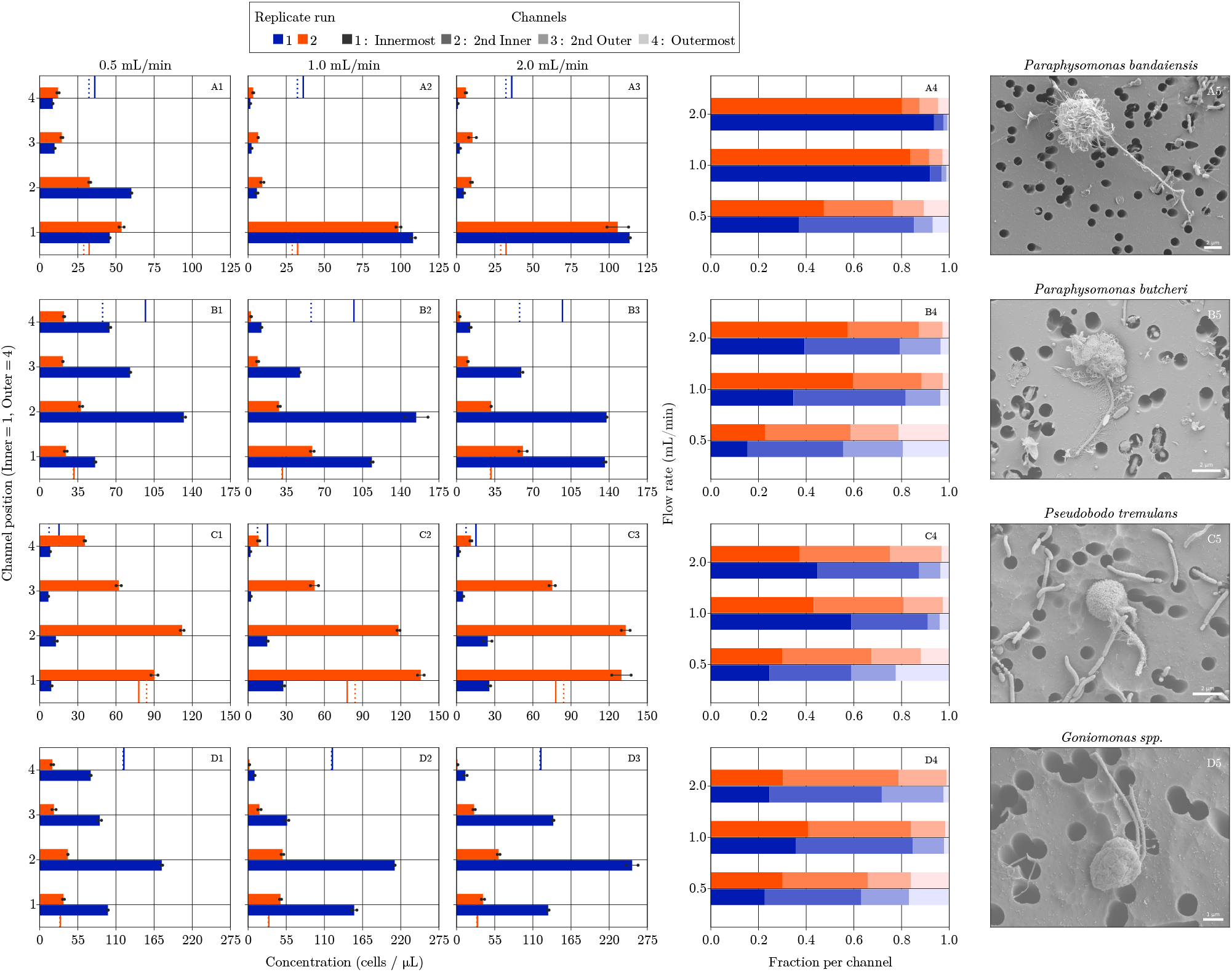
For A) *Paraphysomonas banadaiensis*, B) *Paraphysomonas butcheri*, C) *Pseudobodo tremulans* and D) *Goniomonas* spp., column panels 1-3) show average cell concentration for run 1 (blue bars) and run 2 (orange bars) for each outlet channel from duplicate flow cytometry samples (values indicated by black range bars) for each flow speed. Vertical lines in each plot indicate cell concentration in bulk culture flask before (dashed line) and after (solid line) the transit experiment for each run (indicated by color). Panel 4) Proportion of cell concentration distributed across each channel (shade of color) for each run (color). Panel 5) Scanning electron micrograph of each protist grazer.

### Experiments with microfluidic spiral device

All four nanoflagellates used in this study were able to be concentrated within the range of flow speeds tested (Fig. 2A1-D3). A flow speed of at least 1 mL/min was needed to obtain the highest concentration of cells in the device’s inner two output channels. However, a flow speed of 0.5 mL/min still resulted in different concentrations in each output channel. We did not find a marked increase in concentrating ability when increasing flow speed from 1 mL/min to 2 mL/min. Duplicate experiments for each nanoflagellate culture showed similar results, including the experiments with a physically different device (run 2 for *P. bandaiensis* and *P. tremulans*). We were able to double the cell concentration in at least one output channel compared to the original culture flask for each of these nanoflagellates. The device used in this study was able to handle flow rates up to 2 mL/min without any leaks, however, initial testing with faster rates did result in fluid leaking around the inlet opening as well as occasional bond failure within the device channels.

Output volume of sample fluid through the device was generally not equal among the four output channels. There did not appear to be any pattern regarding which outlet channel contained the most volume; volumes in each channel differed across runs and across nanoflagellate species (see Table S1). Despite volume differences, cells did not appear to be lost in the device or breaking internally. Total cell amount calculated from each output tube volume and output cell concentration was similar to the initial concentration of the sample flask before transiting through the device (Table S2). However, for a small number of sample runs, nanoflagellates appeared to be dividing as the experiment was taking place, determined by the increase in cell concentration in the bulk culture flask before and after the spiral experiment. In these cases, it was difficult to compare the average output amount of cells to the input amount at the beginning and at the end of each experiment.

*Paraphysomonas bandaiensis* was the most affected by the spiral device, as almost all of the cells concentrated in the innermost output channel with a flow speed of at least 1 mL/min. Cell concentration in the innermost channel was over 3 times higher than in the original culture. Results for the three remaining nanoflagellate species were similar to each other, with the majority of the cells moving to the two innermost channels at a flow speed above 1 mL/min. Interestingly, for *Goniomonas* spp., a higher concentration of cells in the inner channels was achieved at 1 mL/min rather than at 2 mL/min.

Bacteria concentrations were also examined in the output channels of the nanoflagellate experiments (Fig. S1). Background bacteria appeared relatively evenly dispersed across all four channels for each speed examined, except for the bacteria found within *P. tremulans* cultures. Bacteria were slightly higher in the outer channel at a low flow speed of 0.5 mL/min, and tended to be slightly concentrated in the two inner channels at higher speeds (Fig. S1). It is unclear if this is due to bacteria clumps or the presence of elongated shapes. Control experiments with a single bacteria strain showed similar results to those for most of the mixed assemblage of bacteria within the nanoflagellate cultures. Cells of *Shimia* strain 13.20C-C appeared fairly evenly distributed across all four outlet channels (Fig. 3). Slight differences in bacteria concentrations among the four output channels were observed, but were not consistent across channels for repeated experiments.

**Figure 3.**
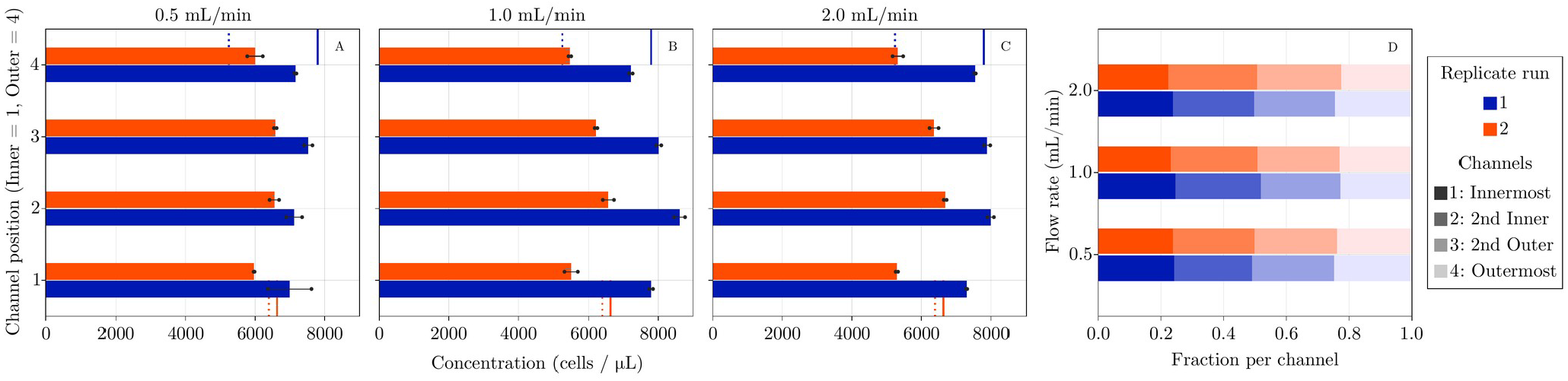
A-C) Average cell concentration for bacteria control strain *Shimia* 13.20C-C for run 1 (blue bars) and run 2 (orange bars) for each outlet channel from duplicate flow cytometry samples (values indicated by black range bars) for each flow speed. Vertical lines in each plot indicate cell concentration in bulk culture flask before (dashed line) and after (solid line) the transit experiment for each run (indicated by color). D) Proportion of cell concentration distributed across each channel (shade of color) for each run (color).

All grazers resumed normal and characteristic motility patterns after transiting through the device. Cultures that were started using transited cells also showed positive growth after a few days of incubation.

## Discussion

We find that a spiral microfluidic device is able to concentrate marine nanoflagellates in the 320 μm size range. To our knowledge, this is the first attempt to use microfluidic devices for the concentration of these cell types. None of the four nanoflagellates showed signs of damage after transiting through the device. We are unaware of other reports of differences in output volume among the outlet channels. These differences in volume, however, do not seem to affect the device’s ability to concentrate cells. Possible reasons for volume differences could be differences in sample viscosity, starting cell concentrations, cell surface stickiness, cell aggregations affecting flow rate, or possible structural imperfections within the PDMS channels.

The degree to which cells were concentrated in a given outlet channel appeared to be size dependent, with larger *P. bandaiensis* cells being most affected by the forces within the device. Concentration of cells did not appear to be affected by motility, extracellular characteristics, or culture density. These observations are consistent with Roth et al. (2018), who found that size was the most important factor determining successful particle separation over particle shape.

Bacteria appeared unaffected by the forces inside the spiral and did not consistently concentrate into a particular output channel. This is likely due to their much smaller size, which is around a micron in diameter. While we did observe elongated forms of bacteria several microns in length (thought to be a defense response to avoid ingestion by predators), such cells also do not appear to be concentrated within the device (Corno and Jürgens 2006). This would indicate that it is the diameter of cell size that is the important dimension rather than the length of the cell. However, more rigorous and defined experiments will be needed to investigate elongated shapes. For the experiments presented here, a cell diameter of three microns appears to be the minimum size for which inertial and drag forces act to move cells into an equilibrium position within the channel.

The spiral microfluidic device enables a quick and effective way to concentrate protists without damage, providing several advantages over traditional centrifugation and membrane filtration approaches. Experiments in this study focus on concentrating nanoflagellate cells and reducing surrounding bacteria concentrations in a sample. For these organisms, such a capability will enable more robust ingestion, growth and other types of experiments. Reduction of background bacteria should also aid in attempts to render cells axenic. This is typically attempted with use of wide-spectrum antibiotics, but these compounds can have a negative and harmful impact on protist cells (Hamilton and Hunter-Cevera unpubl.; Corno and Jürgens 2006; Lowe et al. 2011). Reduction in bacteria numbers and types could allow for a reduced course of antibiotic treatments. These microfluidic devices could also be utilized to size separate protist cells from each other to aid in isolation attempts, especially when additional changes in channel geometry and flow speed are considered. Guan et al. (2013) demonstrated that 15.5 μm diameter beads could be separated from 18.68 μm diameter beads using a spiral device that had trapezoidal channels. The ability to separate such closely sized particles should translate to cell separation capabilities and will provide useful new tools to size select different cell types from the marine environment.

## Comments and Recommendations

The increasing availability of 3D printers with micron printing resolution has greatly reduced the barrier to entry to develop and utilize microfluidic devices for many laboratories. This technology now gives researchers an accessible building platform that requires less time and costly equipment compared with traditional techniques, such as CNC milling. While 3D printing resolution has improved dramatically in recent years, some issues remain for construction of microfluidic devices. We encountered bonding issues between the surface of our casted PDMS and glass slide, which required great precision and care with plasma preparation to ensure a strong bond (+/− 5 seconds plasma exposure would result in bond failure). We hypothesize that these issues were either from an imperfectly smooth surface generated by the 3D printer or possible resin residues leaching into the cured PDMS (Parthiban et al. 2021; Venzac et al. 2021). In comparison, PDMS poured and cured on a petri dish did not demonstrate any bonding issues (e.g. strong, reliable bonds and broad timing range for plasma exposure). We recommend future testing of these parameters during device construction and exploring the use of silanization to create a smooth and non-leaching mold surface (Catterton et al. 2021).

## Conclusions

Marine protist isolation and concentration manipulation has traditionally relied on centrifugation and membrane filtration, which can harm these fragile cell types. We demonstrate the first use of a microfluidic device to concentrate nanoflagellate protists in the 3-20 μm size range. We show that despite irregular cell surfaces, flagella and motility, these cells can be concentrated unharmed in a microfluidic device. While larger nanoflagellates were most affected by the forces within the device, cells as small as 3 μm were still successfully concentrated using flow speeds in the 1-2 mL/min range. We anticipate that such devices will assist researchers in isolation attempts as well as experimental setup requiring concentrated protist cells and reduction in background bacteria levels.

## Acknowledgments

We would like to thank Kasia Hammar and Louis Kerr for assistance with scanning electron microscopy preparation and imaging, Rob Lewis for assistance in repair and tuning of the plasma prep machine, and Emily Stone for assistance with growing bacteria needed for experiments as well as scanning electron microscopy sample preparation. This work was supported by a Hibbitt Early Career Fellowship and Simons Foundation Early Career Investigator Award in Marine Microbial Ecology and Evolution to K. R. Hunter-Cevera.

## Supplementary Figures and Tables

**Figure S1:**
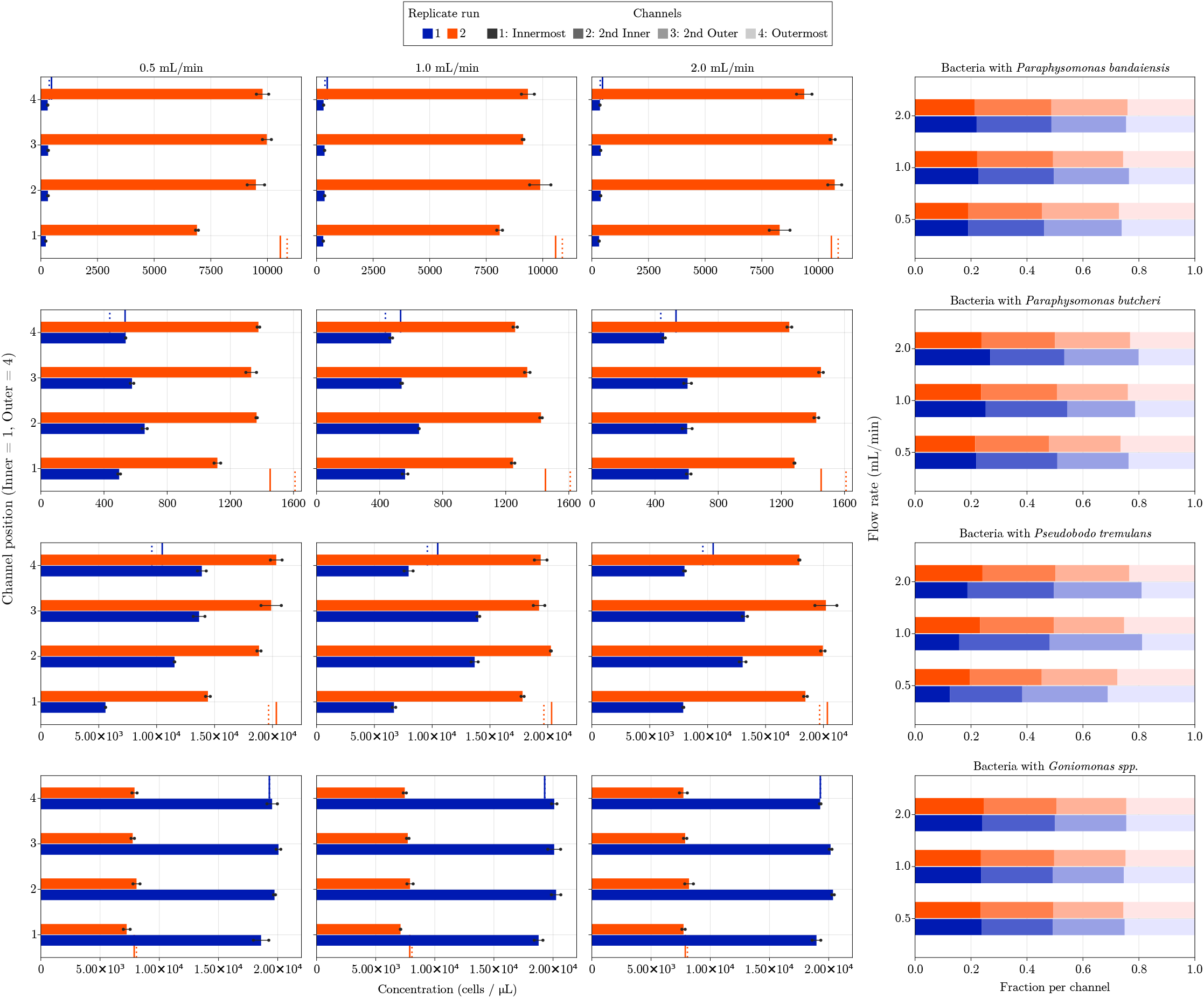
For experiments with A) *Paraphysomonas banadaiensis*, B) *Paraphysomonas butcheri*, C) *Pseudobodo tremulans* and D) *Goniomonas* spp., column panels 1-3) show average background bacteria cell concentration for run 1 (blue bars) and run 2 (orange bars) for each outlet channel from duplicate flow cytometry samples (values indicated by black range bars) for each flow speed. Vertical lines in each plot indicate bacteria cell concentration in bulk culture flask before (dashed line) and after (solid line) the transit experiment for each run (indicated by color). Panel 4) Proportion of bacteria cell concentration distributed across each channel (shade of color) for each run (color).

**Table S1:**
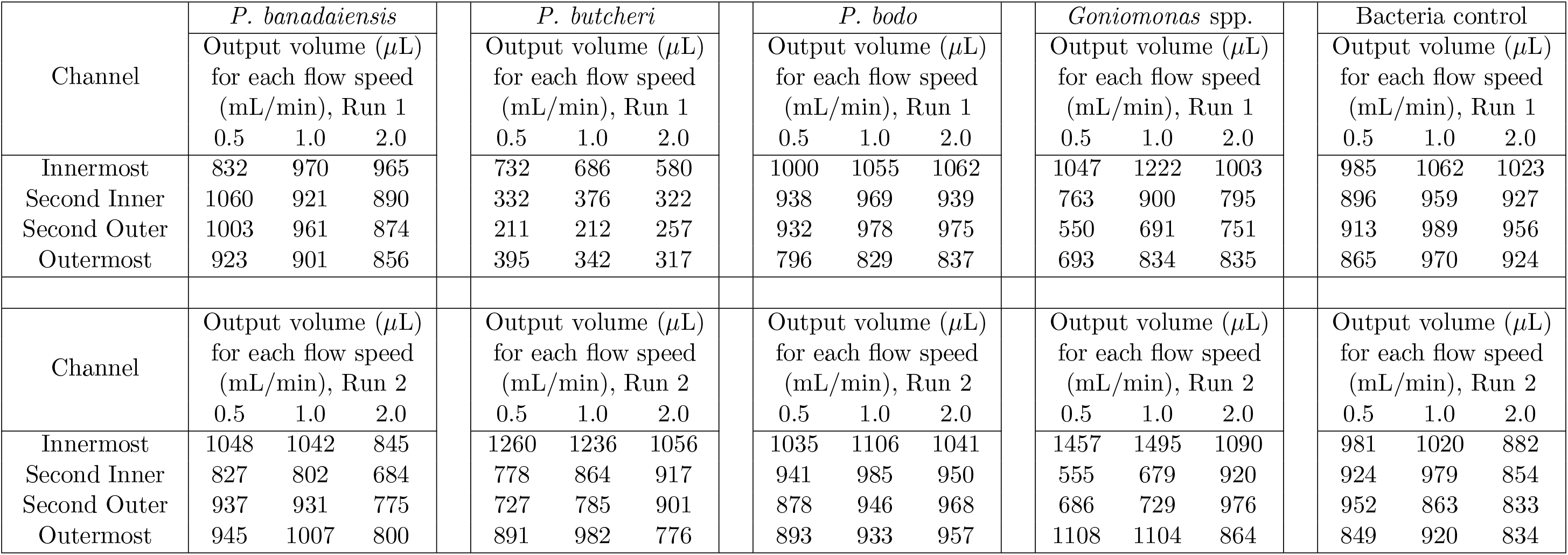
Output volumes for each outlet channel for each experiment. In general, outlet volumes were similar across channels, but for some experiments, varied up to 50% across outlets.

**Table S2:** Average cell concentration (cells/*μ*L) across all four outlet channels for each experiment and cell type. Note this is not an average of values presented in Fig. 2. Rather, average concentration is calculated by determining total number of cells in each outlet (outlet volume multiplied by outlet cell concentration), divided by total combined outlet volume (i.e. sum of all outlet channel volumes). Concentrations listed for before and after each experiment reference bulk culture flask values.

| Culture/Exp. | Before Exp. | 0.5 mL/min | 1.0 mL/min | 2.0 mL/min | After Exp. |
| --- | --- | --- | --- | --- | --- |
| *P. bandaiensis*, Run 1,<br>Avg. protist conc. | 32.33 | 31.37 | 30.29 | 32.41 | 36.14 |
| *P. bandaiensis*, Run 1,<br>Avg. bacteria conc. | 371.47 | 292.57 | 328.93 | 365.19 | 471.03 |
| *P. bandaiensis*, Run 2,<br>Avg. protist conc. | 28.89 | 28.89 | 31.63 | 35.07 | 32.43 |
| *P. bandaiensis*, Run 2,<br>Avg. bacteria conc. | 10868.11 | 8960.34 | 9062.4 | 9687.24 | 10568.72 |
| *P. butcheri*, Run 1,<br>Avg. protist conc. | 57.8 | 74.29 | 92.94 | 96.47 | 97.09 |
| *P. butcheri*, Run 1,<br>Avg. bacteria conc. | 436.62 | 548.14 | 560.47 | 576.7 | 533.07 |
| *P. butcheri*, Run 2,<br>Avg. protist conc. | 31.05 | 26.09 | 27.64 | 28.79 | 31.49 |
| *P. butcheri*, Run 2,<br>Avg. bacteria conc. | 1609.49 | 1276.59 | 1307.85 | 1352.43 | 1452.06 |
| *P. tremulans*, Run 1,<br>Avg. protist conc. | 7.43 | 9.35 | 12.44 | 14.99 | 15.24 |
| *P. tremulans*, Run 1,<br>Avg. bacteria conc. | 9582.97 | 10967.95 | 10596.71 | 10539.1 | 10480.81 |
| *P. tremulans*, Run 2,<br>Avg. protist conc. | 84.12 | 76.15 | 81.57 | 88.06 | 77.94 |
| *P. tremulans*, Run 2,<br>Avg. bacteria conc. | 19664.29 | 18219.9 | 19135.98 | 19116.55 | 20335.24 |
| *Goniomonas spp.*, Run 1,<br>Avg. protist conc. | 120.41 | 110.14 | 116.20 | 132.65 | 121.49 |
| *Goniomonas spp.*, Run 1,<br>Avg. bacteria conc. | 19299.32 | 19353.87 | 19697.08 | 19635.36 | 19290.54 |
| *Goniomonas spp.*, Run 2,<br>Avg. protist conc. | 29.26 | 28.18 | 29.35 | 31.99 | 29.93 |
| *Goniomonas spp.*, Run 2,<br>Avg. bacteria conc. | 8067.03 | 7655.33 | 7448.87 | 7885.69 | 7881.42 |
| *Shimia* strain 13.20C-C,<br>Run 1, Avg. conc. | 5243.72 | 7196.86 | 7906.23 | 7674.08 | 7795.64 |
| *Shimia* strain 13.20C-C,<br>Run 2, Avg. conc. | 6393.75 | 6277.16 | 5927.99 | 5912.00 | 6630.91 |

